# PlasmoRhythm: A comprehensive database for malaria-associated rhythmic transcripts, proteins, and metabolites

**DOI:** 10.64898/2026.10.11.758397

**Authors:** Sourbh Rankawat, Kavita Kundal, Subashani Singh, Sandip Das, Bianca Parisi, Bhanu Teja Korra, Filipa Rijo-Ferreira, Rahul Kumar, Sandipan Ray

## Abstract

**Motivation:** The malaria transmission cycle is temporally organized across three interacting biological systems: the parasite, vertebrate host, and mosquito vector. Rhythms in parasite development, host physiology, and vector behavior govern infection dynamics, transmission, and the efficacy of stage-specific antimalarial interventions. However, the molecular architecture connecting these rhythms remains difficult to interrogate because time-resolved omics datasets are distributed across species, experimental paradigms, and analytical frameworks, with no unified resource for comparative analysis.

**Results:** We developed PlasmoRhythm, a curated database integrating 67 transcriptomic, proteomic, and metabolomic datasets from 15 studies, encompassing 1,126 sampling events across *Plasmodium* species, mammalian hosts, and *Anopheles* vectors. Harmonized metadata and rhythmicity outputs from JTK_CYCLE, ARSER, Lomb–Scargle, and the meta2d function support interactive cross-study interrogation of molecular profiles. Dedicated modules enable the exploration of rhythmic virulence factors and antimalarial drug interactors, providing a unified framework for hypothesis generation across the malaria transmission cycle.

**Availability:** PlasmoRhythm is freely accessible at https://project.iith.ac.in/cgntlab/PlasmoRhythm/.

## 1 INTRODUCTION

Malaria, caused by protozoan parasites of the genus *Plasmodium*, remains a leading cause of morbidity and mortality worldwide, sustained by a complex lifecycle alternating between human and mosquito hosts (World malaria report 2025). The transmission cycle begins when an infected *Anopheles* mosquito inoculates sporozoites into the human bloodstream, initiating an asymptomatic replication phase within hepatocytes. Upon completion of this hepatic stage, merozoites are released into the circulation, where they invade erythrocytes and initiate the intraerythrocytic developmental cycle (IDC), a repeating process of growth, replication, and egress (Cowman *et al*. 2016). The IDC follows a species-specific periodicity, usually in multiples of ∼24 hours. This periodicity drives malaria’s rhythmic fevers, providing possible evidence for an intrinsic timekeeping mechanism within the parasite (Hawking 1970, Garcia, Markus, and Madeira 2001, Sadykov and Rijo-Ferreira 2020). Recent transcriptomic studies have demonstrated that the IDC is supported by widespread rhythmic gene expression, with 87–92% of the *Plasmodium falciparum* transcriptome exhibiting oscillatory behavior, with a periodicity of approximately 48 h *in vitro*, independent of host cues (Smith *et al*. 2020). Similarly, approximately 80% of genes in *Plasmodium chabaudi* exhibit a ∼24 h periodicity, even when the host is kept in constant darkness (Rijo-Ferreira *et al*. 2020). Although little is known about the genetic basis of parasite rhythms, some correlational studies have identified certain genes associated with rhythmic behavior. For example, disruption of the expression of the serpentine receptor Sr10 in *P. chabaudi* led to a reduced number of cycling genes, potentially linking G-protein-coupled receptor (GPCR) signaling to the rhythmicity of malaria during its intraerythrocytic stage (Subudhi *et al*. 2020).

Beyond intrinsic clocks in *Plasmodium*, much of the synchronized oscillations observed align with the rhythms of its host or vector (Rijo-Ferreira *et al*. 2020, Smith *et al*. 2020, O’Donnell, Prior, and Reece, 2020). For example, a recent study found that in *Plasmodium vivax*-infected individuals, parasite development is phase-aligned with the host’s circadian transcriptome, potentially facilitating immune evasion or metabolic synchronization to host metabolite availability (Motta *et al*. 2023). Notably, ∼50% of the host immune factors thought to be important for controlling malaria infection also exhibit daily oscillations (Rankawat *et al*. 2023), and altering the host’s circadian environment affects parasite development and rhythmicity during the intraerythrocytic cycle (Rijo-Ferreira *et al*. 2020, Hirako, Ramalho, and Gazzinelli 2025). hythms in the malaria vector have also been recently well classified. Studies on *Anopheles gambiae* have revealed rhythmic biting patterns and tissue-specific transcriptional oscillations in the mosquito head and body, while work on *Anopheles stephensi* has shown extensive rhythmic gene expression in the salivary glands, where parasites are harbored before transmission, under both light-dark and constant dark conditions (Rund *et al*. 2011, Bento *et al*. 2025).

The efficacy of most antimalarial drugs often depends on the parasite’s developmental stage (Wilson *et al*. 2013). Elucidating how host and parasite rhythms co-depend could be vital for better understanding the optimal timing of administering anti-malarial therapies, since pharmacokinetic and pharmacodynamic processes are governed by robust 24 h rhythms (Ruben *et al*. 2019, Dong *et al*. 2020).

Altogether, decoding the rhythms of parasites, hosts, and vectors is crucial for understanding malaria pathogenesis, dynamic host-pathogen interactions, and the potential time dependence of antimalarial interventions. Despite the increasing number of publications featuring large-scale, systems-level time-series datasets on malaria parasites, hosts, and vectors (**Figure 1**), there is a genuine need for an integrated web-based platform that enables comprehensive exploration, analysis, and integration of all rhythms involved in malaria pathogenesis and infection.

**Figure 1.**
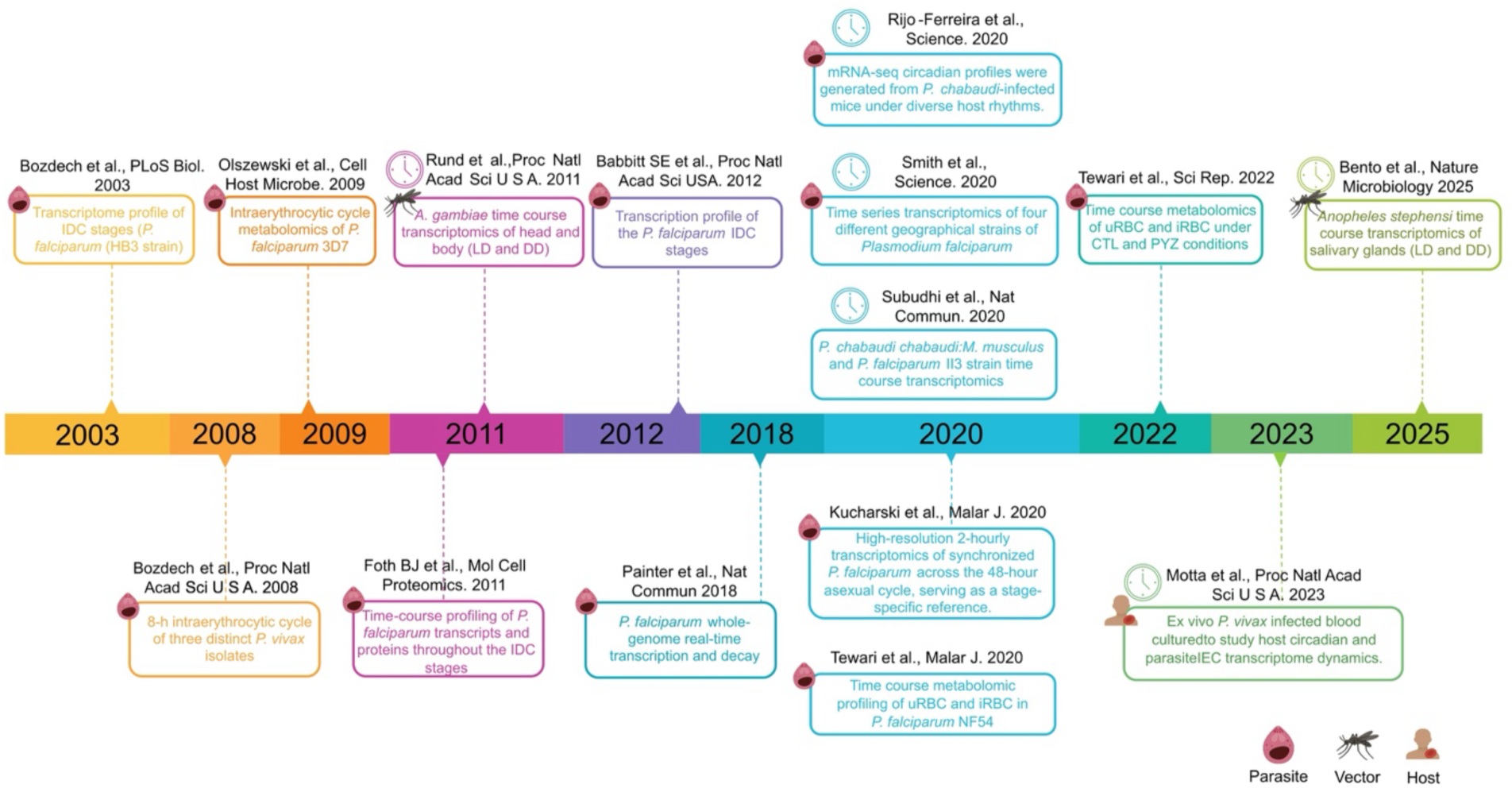
Timeline of representative time-series studies in malaria research. This timeline presents landmark time-series studies at various omics levels on Plasmodium parasites, mosquito vectors, and host systems from 2000 to 2025. Each entry indicates the publication, organism(s), and study focus, with icons for parasite, vector, and host systems. The timeline highlights advancements in the high-resolution transcriptomic, proteomic, and metabolomic profiling, including studies of the intraerythrocytic developmental cycle (IDC) and cross-species analyses. Studies marked with a clock icon specifically address circadian or diel rhythmicity. This figure illustrates the evolution and expansion of temporal omics research in malaria, culminating in the integrated datasets curated within the PlasmoRhythm database.

To address this gap, we developed the PlasmoRhythm database, the first comprehensive web-based platform dedicated to capturing genome-wide oscillatory dynamics across the malaria transmission cycle **(Figure 2)**. By integrating several publicly available transcriptomic, proteomic, and metabolomic time-series data from *Plasmodium* species, mosquito vectors, and host systems, PlasmoRhythm offers a valuable platform for systematic analysis of the biological rhythms of malaria-associated species. In PlasmoRhythm, molecular rhythmicity is assessed using multiple well-established algorithms, with customizable thresholds, interactive plots, and advanced search tools to support examinations of time-dependent biological processes in malaria. PlasmoRhythm primarily focuses on three imperative aspects of clock-infection biology in malaria: (a) interactive visualization of the time-dependent expression or abundance of transcripts, proteins, and metabolites under diverse conditions and species, (b) rhythmic regulations of putative malaria virulence factors and cross-system exploration of parasite-host-vector rhythmic coupling to a clock-informed transmission biology and therapeutic interventions, and (c) identifying candidate targets for circadian medicine (chronotherapy) for malaria by mapping rhythms in antimalarial targets and druggable genes relevant to pharmacokinetic windows (such as a short plasma half-life). Overall, PlasmoRhythm provides a valuable and unique resource to the malaria, vector, and circadian research communities for in-depth study of clock-infection biology in malaria.

**Figure 2.**
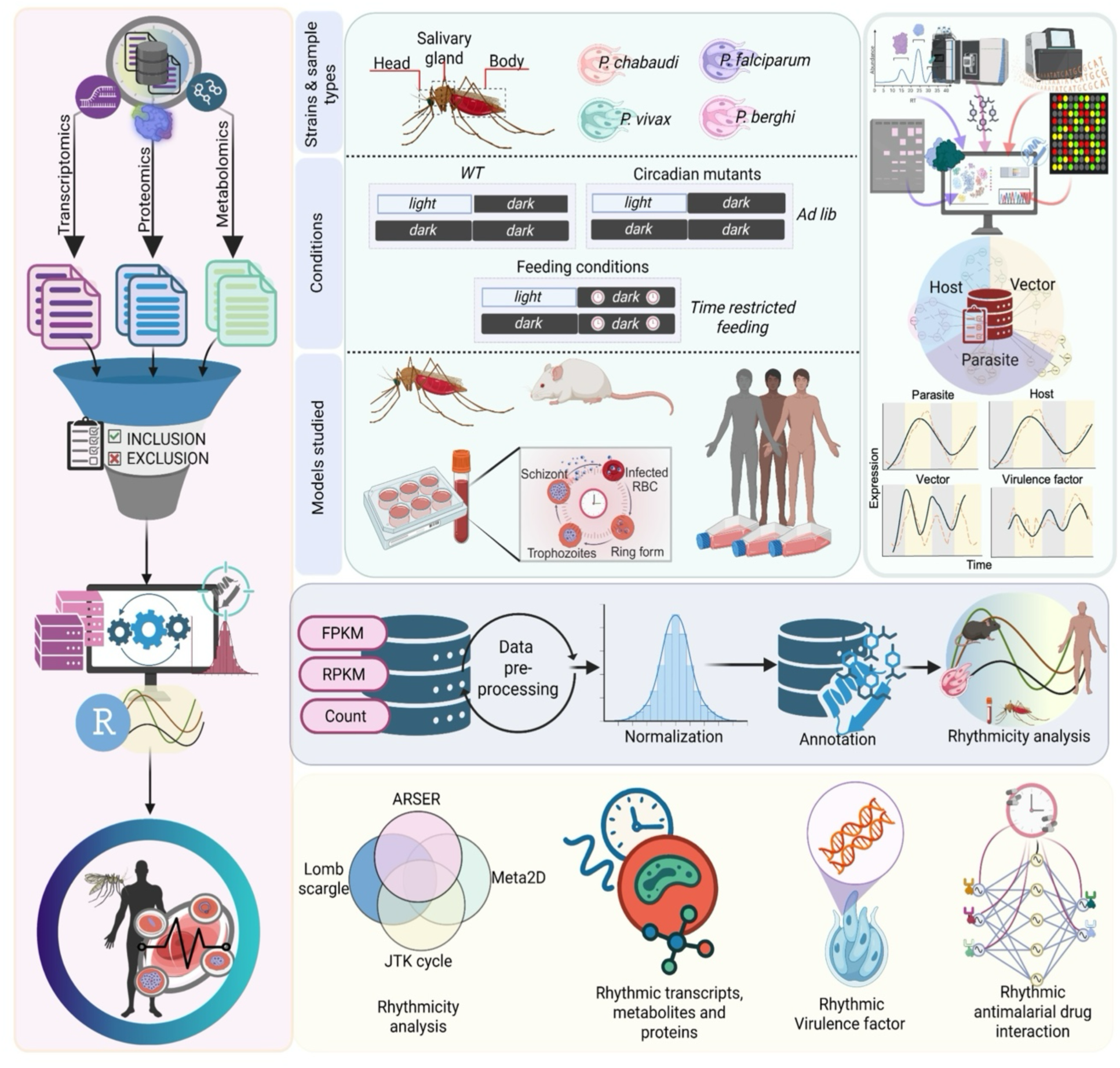
Construction and content of PlasmoRhythm: a comprehensive database for malaria-associated rhythmic transcripts, proteins, and metabolites. The PlasmoRhythm database (https://project.iith.ac.in/cgntlab/PlasmoRhythm/) is designed to provide an interactive web-based platform for comprehensive analyses of the rhythmic patterns in malaria parasites, their hosts, and mosquito vectors. It integrates several transcriptomic, proteomic, and metabolomic datasets from multiple Plasmodium species and Anopheles mosquitoes under different experimental conditions and strains. Users can input gene IDs, protein IDs, or metabolite names and visualize their rhythmic profiles. Additionally, they can adjust the statistical parameters using multiple integrated algorithms. The platform generates interactive visualizations of phase, amplitude, and period of rhythmic patterns, along with insights into parasite virulence factors and interactions with antimalarial drugs.

## 2 METHODS

### 2.1 Dataset collection and curation

We systematically searched publicly available time-course datasets related to malaria parasites, their hosts, and vectors using PlasmoDB (https://plasmodb.org/plasmo/app (Aurrecoechea *et al*. 2009, Alvarez-Jarreta *et al*. 2024), PubMed (https://pubmed.ncbi.nlm.nih.gov/), and Google Scholar (https://scholar.google.com/). To identify relevant studies, we used combinations of keywords including “*Plasmodium*,” “malaria,” “time-course,” “circadian,” “rhythm,” “transcriptome,” “expression profiling,” “parasite stage,” “vector,” and “host.”

We applied stringent inclusion criteria across different omics layers to ensure robust rhythm detection and sufficient temporal resolution. Transcriptomic datasets require sampling intervals of ≤ 6 h and ≥ 9 time points to capture ≥ 2 complete 24 h cycles for reliable oscillation detection, consistent with published guidelines for genome-scale rhythmicity analysis (Hughes *et al*. 2017). Three *ex vivo P. vivax* datasets (6-8 in **Table S1**) were exceptions to the criteria. They had only nine irregularly spaced measurements spanning 9-43 h of the IDC, but they represent rare longitudinal transcriptomic profiles from clinical *P. vivax* isolates and cover most of the developmental cycle.

We included metabolomics datasets with ≤8 h intervals, given the current paucity of high-sampling resolution metabolomics datasets in malaria research. At this resolution, the available datasets comprise 7 timepoints sampled across the full 48-h IDC (0–48 h at 8-h intervals), capturing one complete IDC cycle; this is less stringent than the transcriptomic criterion in terms of cycle coverage but represents the maximum temporal resolution datasets currently available for metabolome-level time-series analysis of malaria parasites.

Proteomic datasets were included if they had a resolution of ≤ 6 h, ≥ 4 time points/day, and encompassed both single-cell proteomics (e.g., *A. stephensi* salivary glands) and 2D-DIGE profiles coupled with mass spectrometry (e.g., *P. falciparum* IDC). These criteria were selected to balance the requirements of rhythmicity detection with the limited availability of high-resolution proteomic time-series data in malaria parasites. Unlike transcriptomic datasets, available time-series proteomic studies in malaria typically entail greater analytical complexity, lower coverage, and lower sampling resolution, necessitating inclusion criteria that preserve statistical interpretability while maximizing biological representation.

Employing the above-mentioned section criteria, we defined 67 high-quality time-series datasets from 15 studies comprising a total of 1,126 data points. These included foundational work on *P. falciparum and P. vivax* IDC (Bozdech *et al*. 2003, 2008), host-parasite synchronization in *P. vivax* infections (Motta *et al*. 2023), vector rhythm studies (Rund *et al*. 2011, Bento *et al*. 2025), transcriptome-wide analyses of parasite developmental rhythms (Rijo-Ferreira *et al*. 2020, Smith *et al*. 2020, Subudhi *et al*. 2020), global transcriptional profiling of *P. falciparum* blood-stage development (Olszewski *et al*. 2009, Babbitt *et al*. 2012, Painter *et al*. 2018), proteomic characterization of parasite cell cycle progression (Foth *et al*. 2011), metabolic regulation during infection (Kucharski *et al*. 2020), host cell remodeling (Tewari *et al*. 2020), and erythrocyte invasion dynamics (Tewari *et al*. 2022) (**Table S1**).

### 2.2 Data preprocessing

Raw transcriptomic datasets (n = 55), comprising 15 microarray and 40 RNA-Seq datasets, were sourced from various platforms, each requiring specific preprocessing to ensure consistency across datasets. Wherever possible, we adhered to the preprocessing pipelines outlined by the original authors to maintain fidelity to the original study designs. Some datasets were used directly as provided in the original studies without further preprocessing (**Table S1**). Microarray datasets were downloaded in “gpr” format and pre-processed using the R statistical platform. Background correction was performed using the “normexp” function, and quantile normalization was performed using the “normalizeBetweenArrays” function.

All normalized intensities were mapped to gene identifiers as follows: *Plasmodium* gene IDs were mapped using PlasmoDB release 64 (Aurrecoechea *et al*. 2009); to ensure uniformity, *P. vivax* Sal-1 clinical isolate IDs were also mapped via PlasmoDB to the *P. vivax* P01 reference genome identifiers. Vector annotations (*A. gambiae and A. stephensi*) using VEuPathDB (https://veupathdb.org) (Alvarez-Jarreta *et al*. 2024), and *H. sapiens* gene IDs using SynGO portal (https://syngoportal.org) (Koopmans *et al*. 2019), which is integrated with the Gene Ontology resource. Compound names for metabolites were uploaded to MetaboAnalyst 6.0 (https://www.metaboanalyst.ca) (Pang *et al*. 2024), and the ‘Compound ID Conversion*’* module was used to retrieve identifiers from the Kyoto Encyclopedia of Genes and Genomes (KEGG), the Human Metabolome Database (HMDB), and PubChem.

Time-series RNA-seq datasets generated using Illumina sequencing platforms were provided in normalized RPKM/FPKM format and directly incorporated into the analysis. To remove lowly expressed genes, a threshold cutoff of 0.5 was applied, and genes with expression values < 0.5 in more than 30% of samples were excluded from further analysis (Kucharski *et al*. 2020, Rijo-Ferreira *et al*. 2020, Smith *et al*. 2020). RNA-seq preprocessing and expression-filtering procedures were selected based on the expression measure and characteristics of each dataset (**Table S1**). The dataset (Subudhi *et al*. 2020), available as raw counts, was normalized using TMM (trimmed mean of M-values) via the “gdcVoomNormalization” function in R. A dataset from (Motta *et al*. 2023), also in raw counts, was processed with quantile normalization before further analyses. We processed the metabolomics (Olszewski *et al*. 2009, Tewari *et al*. 2020, 2022) and proteomics (Foth *et al*. 2011, Bento *et al*. 2025) datasets as described in the original publications (**Table S1**).

### 2.3 Identification and annotation of *Plasmodium* virulence factors

The putative virulence factors in *P. falciparum* and *P. vivax* were curated through an extensive literature review using PubMed. Search terms included “*Plasmodium* virulence factor,” “erythrocyte invasion,” “antigenic variation,” “immune evasion,” and “cytoadherence,” along with species-specific terms. Studies published between 2000 and 2025 were reviewed, and genes were selected based on experimental evidence (e.g., knockouts, protein-binding assays), conservation across *Plasmodium* species, and homology with virulence proteins in other apicomplexans.

Pseudogenes and truncated sequences were excluded using PlasmoDB’s “bona fide” filters and manual review. Based on published literature, genes were classified by function (e.g., erythrocyte invasion, antigenic variation, immune evasion). Gene identifiers, functional annotations, and Gene Ontology (GO) terms were obtained from PlasmoDB release 64 (Aurrecoechea *et al*. 2009).

### 2.4 Cataloguing interactors of antimalarial drugs

Commonly used FDA-approved antimalarial drugs were retrieved from DrugBank (release 5.1.13; CC BY-NC 4.0, https://go.drugbank.com/) (Knox *et al*. 2024). To facilitate pharmacokinetic classification, drugs were grouped based on their plasma half-lives in adults: short (≤15 h), medium (15–24 h), long (1–14 days), and ultra-long (>14 days).

Drug–target interactions were identified using the STITCH 5 (http://stitch.embl.de/) (Szklarczyk *et al*. 2016) via the stringApp plugin in Cytoscape (v3.10.3) (Doncheva *et al*. 2023). The STITCH database integrates multiple lines of evidence, including experimental data, curated databases, co-expression, gene fusion, co-occurrence, and text mining (Szklarczyk *et al*. 2016). Queries were performed at the species level for *H. sapiens*, *P. falciparum*, *P. vivax*, and *P. chabaudi*. All drug–target interactions meeting the STITCH combined-confidence threshold (≥0.400) were retained rather than being restricted to a fixed top-N set. The interface displays the combined confidence score and evidence channels and allows users to sort or filter interactions by confidence category, from lower to higher confidence; disconnected nodes were excluded.

To supplement database predictions, we performed a manual literature review focusing on experimentally validated drug–target interactions apart from those captured by STITCH. PubMed and Google Scholar were searched using terms such as “antimalarial drug target validation,” “*Plasmodium* drug interaction,” and with specific names of the compounds - Artemether, Artenimol, Primaquine, Quinidine, Amodiaquine, Methylene blue, Quinine, Proguanil, Atovaquone, Lumefantrine, Pyrimethamine, Piperaquine, Halofantrine, Tafenoquine, Mefloquine, Chloroquine, and Hydroxychloroquine.

Artemisinin and its derivatives were subjected to additional manual curation because their mode of action involves extensive protein alkylation and multi-target interactions that are not comprehensively represented in interaction databases. To ensure broad coverage of experimentally supported artemisinin targets, we integrated evidence from multiple chemical proteomics and biochemical studies. These included an alkyne-tagged artemisinin probe study employing click-chemistry enrichment and LC-MS/MS (Wang *et al*. 2015), activity-based protein profiling studies using artemisinin-derived probes (Ismail *et al*. 2016), a photoaffinity-based activity-based protein profiling study (Gao *et al*. 2024), and targeted biochemical analyses demonstrating direct binding and inhibition of PfDdi1 by artemisinin (Onchieku *et al*. 2021). Orthologous targets in *Plasmodium vivax* and *Plasmodium chabaudi* were retrieved using PlasmoDB.

### 2.5 Rhythmicity analysis

Rhythmic expression analysis was performed using the MetaCycle R package, which integrates three algorithms, JTK_CYCLE, Lomb-Scargle (LS), and ARSER, via the meta2d function (Wu *et al*. 2016). JTK_CYCLE is a non-parametric, rank-based method that compares expression profiles to cosine curves (Hughes, Hogenesch, and Kornacker 2010). The Lomb-Scargle method is a spectral analysis algorithm optimized for unevenly sampled data (Scargle 1982). ARSER employs adaptive harmonic regression to improve periodicity detection (Yang and Su 2010). The default setting of the *meta2d* function, Fisher’s method, was used to combine the *p*-values from the three algorithms it integrates. Period, phase, and amplitude were estimated for each feature. We implemented only the Lomb-Scargle method for rhythmicity analysis for certain datasets with irregular sampling intervals or missing data points (Bozdech et al. 2003, 2008, Babbitt et al. 2012), as other periodicity algorithms (ARSER, JTK_CYCLE, and meta2d) were found unsuitable for these datasets.

MetaCycle parameters (ARSdefaultPer, minper, and maxper) were selected based on the known biological periodicity and the temporal resolution of each dataset. Circadian datasets were analyzed using period ranges centered on ∼24 h, whereas *Plasmodium* IDC datasets were analyzed using period ranges centered on ∼48 h. Complete dataset-specific parameter settings and details on the availability of rhythmicity detection algorithms for each dataset are provided in Supplementary **Table S1**.

To assess the robustness of rhythmicity detection and estimate false-positive rates, permutation-based benchmarking was performed on representative transcriptomic datasets spanning distinct biological and technical contexts (Datasets 24, 19). Timepoint labels were randomly permuted (n = 20) to disrupt temporal structure prior to rhythmicity analysis, using the same MetaCycle workflow and parameter settings as for the original data. Rhythmic candidate counts obtained from the original and permuted datasets were compared using both raw *p*-values and Benjamini–Hochberg (BH)-adjusted *q-*values (**Table S1**). After BH adjustment, the randomly permuted data yielded almost null rhythmic entries in ARSER and JTK_CYCLE (<0.01% of the original calls), but meta2d retained a few rhythmic entries (0.7% and 1.5% of the original calls in Datasets 19 and 24, respectively).

Raw p-values and Benjamini–Hochberg (BH)-adjusted (*q*)-values were calculated for each dataset. All MetaCycle outputs, including period, phase, amplitude, raw *p*-values, and BH-adjusted *q-*values, were retained in the database without imposing a fixed rhythmic/non-rhythmic classification. Unless otherwise stated, illustrative analyses presented in this manuscript used ARSER-derived BH-adjusted *q*-values < 0.05.

### 2.6 Construction of the database

The PlasmoRhythm web platform was developed using a LAMP stack (Linux, Apache, MySQL, PHP), with Java modules supporting data handling and preprocessing. Front-end interactions are built with HTML, CSS, and JavaScript, with dynamic visualizations rendered using the Canvas.js and Cytoscape.js libraries. The platform is hosted on an Apache server maintained at the Indian Institute of Technology Hyderabad. The database implementation described in this manuscript corresponds to PlasmoRhythm version 1.0, released in February 2026.

## 3 RESULTS

### 3.1 Data integration overview

We integrated 67 high-resolution time-series datasets spanning over two decades of malaria research, encompassing multi-omics time-series data from *Plasmodium* parasites, mosquito vectors, and mammalian hosts **(Figure 1)**. The database incorporates 55 transcriptome datasets comprising 81,068 transcripts measured across 1,028 time points, from major *Plasmodium* species and strains including *P. falciparum* (3D7, D6, SA250, II3, FVO-NIH, HB3, Dd2 and 3D7attB strains/clones), *P. vivax* (P01 and three SMRU clinical isolates), *P. chabaudi chabaudi*, *P. berghei*, from two mosquito *Anopheles* species, *A. stephensi* and *A. gambiae*, and human hosts, *Homo sapiens*. Proteomics data include 2,768 proteins from *A. stephensi* and *P. falciparum* (Dd2 strain) quantified across 28 time points, while metabolomics data comprise 3,160 metabolites from 10 datasets spanning 70 time points. The timeline of circadian-omics studies in malaria reveals a steady increase in both temporal resolution and multi-omic depth. As expected by its robust quantitative power, transcriptomic datasets currently dominate (∼80%), with proteomics and metabolomics providing complementary molecular layers.

PlasmoRhythm unifies several high-throughput omics-level datasets, representing 1,126 unique sampling events (data points), into a single platform that supports cross-species and cross-condition comparisons in malaria. The collection encompasses all systems in which *Plasmodium* can be manipulated and tracked, including *in vitro* cultures, animal systems, and clinical samples, under various experimental conditions, such as different parasite strains, feeding regimens, and light cycles. **Figure 2** provides an overview of the PlasmoRhythm database architecture and data processing pipeline. The system architecture includes standardized normalization procedures, annotation protocols, and defined inclusion and exclusion criteria for time-series transcriptomic, proteomic, and metabolomic datasets. To ensure robust rhythmicity detection, all datasets undergo consistent preprocessing and analysis using multiple validated algorithms (JTK_Cycle, ARSER, Lomb-Scargle, and meta2d; see Methods). The platform features interactive tools to visualize rhythmic patterns using user-friendly graphs. Additionally, curated annotations of virulence gene families in malaria and FDA-approved antimalarial drug targets are incorporated, along with their oscillation profiles and rhythmicity parameters, enabling the targeted exploration of molecular rhythms in host, parasite, and vector across different conditions.

### 3.2 Taxonomic and biological composition

In PlasmoRhythm, the majority of data originates from *Plasmodium* parasites (74.62%), with complementary datasets from the infected human hosts (*H. sapiens*) (14.94%) and mosquito vectors (10.44%) **(Figure 3A)**. Among malaria parasites, *P. falciparum* is most represented (50.0%), followed by *P. vivax* (24.0%), *P. chabaudi* (22.0%), and *P. berghei* (4.0%). While malaria mosquito vector datasets include *A. gambiae* (5.97%) and *A. stephensi* (4.47%), with transcriptional profiling focused on tissues relevant to transmission, such as the salivary glands and head-body segments.

**Figure 3.**
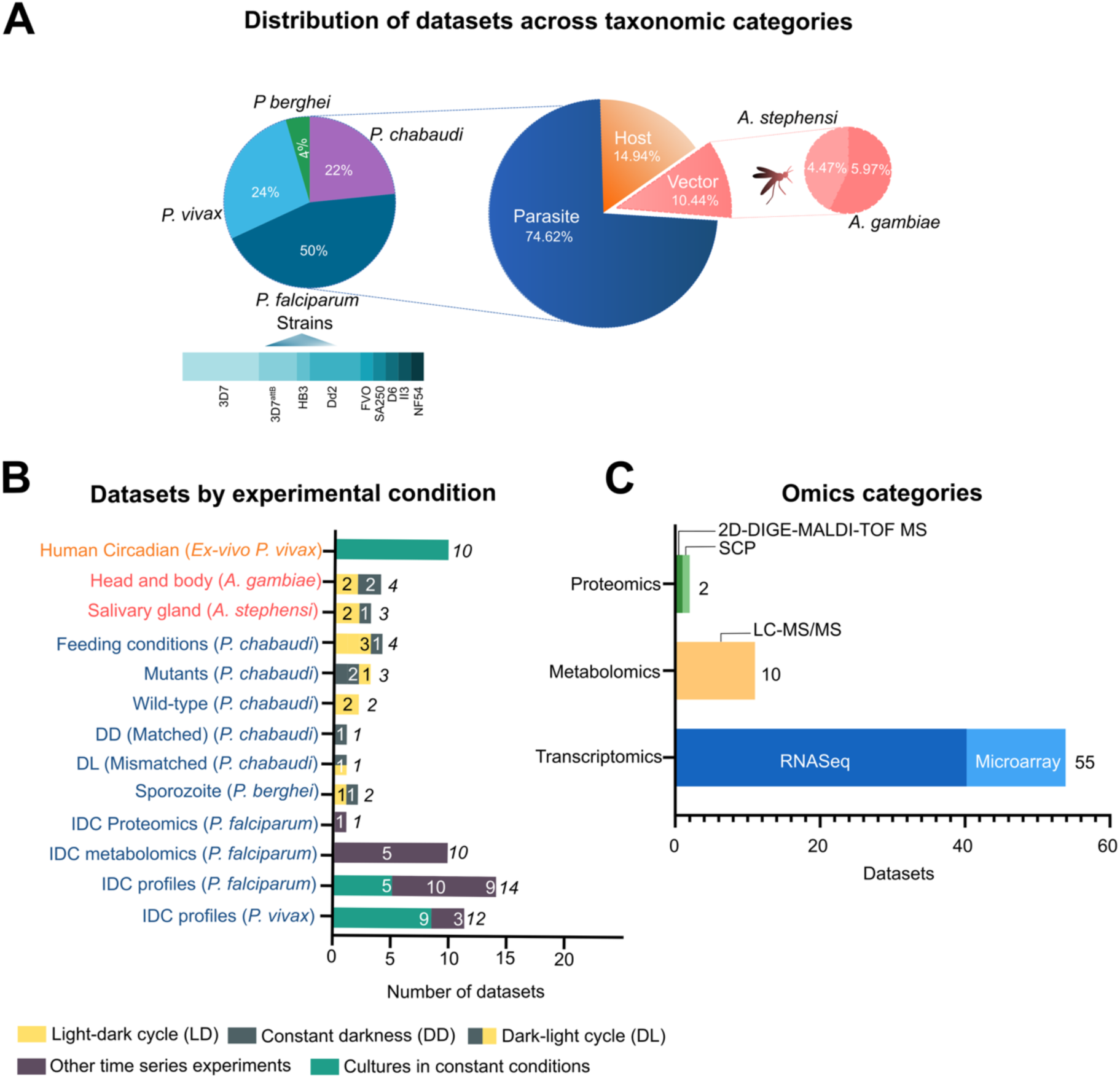
Overview of the taxonomic, experimental, and omics diversity curated within the PlasmoRhythm database. **(A)** Distribution of datasets across biological categories, showing the proportion of datasets derived from parasites (blue), hosts (orange), and mosquito vectors (red). The smaller pie charts further define the parasite and vector datasets by species. **(B)** Discrimination of datasets classified by experimental conditions. Bars indicate dataset counts for each condition across different biological systems, grouped by taxonomic category. The y-axis labels are color-coded to correspond to the broad taxonomic categories defined in Panel A (Host: orange; Vector: red; Parasite: blue). Bar colors represent the specific experimental conditions (e.g., light-dark cycling, constant darkness, or constant environmental conditions), as indicated by the key at the bottom**. (C)** Distribution of datasets by omics category, highlighting the predominance of transcriptomics (blue: RNA-seq; light blue: microarray), with additional representation from metabolomics (orange) and proteomics [green: two-dimensional fluorescence difference gel electrophoresis (2D-DIGE) coupled with matrix-assisted laser desorption ionization–time of flight mass spectrometry (MALDI-TOF MS), and single-cell proteomics (SCP)].

### 3.3 Experimental conditions

All datasets were classified according to different experimental paradigms (**Figure 3B, Table S1**): **Light-regime animal studies:** These primarily include photoperiod-controlled experiments (e.g., LD (light-dark), DD (dark-dark), DL (dark-light) cycles) and feature 11 datasets on *P. chabaudi* infection in mice. These datasets capture how external cues (light cycles and feeding schedules) were applied to hosts and vectors, which may entrain parasite rhythms and influence synchronization.

Four datasets involve wild-type mouse infections under different photoperiods: two standard LD (12 h light:12 h dark), one DL condition where the host rhythm is shifted by 12 h relative to the parasite, and one under constant darkness (DD). Four datasets vary in feeding schedules (night-restricted, ad libitum, or distributed feeding) under LD or DD conditions. In time-restricted feeding experiments, food access was limited to a defined phase of the 24-h cycle, allowing separation of feeding-driven signals from light-driven entrainment. Three additional perturbation datasets include clock-deficient hosts (*FBXL3* KO and *CRY1/CRY2* KO under DD) and a parasite serpentine-receptor knockout (*PchSR10* under LD). Together, these 11 datasets provide a comprehensive framework for analyzing circadian synchronization and the fitness consequences of its disruption during malaria infection.

We also incorporated *P. berghei* sporozoite transcriptomes under LD and DD, and seven mosquito datasets spanning both species: *A. stephensi* salivary gland transcriptomes (LD/DD) and a salivary gland day–night proteome, as well as *A. gambiae* head- and body-segment transcriptomes across LD and DD. These datasets enable assessment of light-dependent rhythmicity across the parasite life cycle and mosquito vector tissues.

### 3.4 Constant-condition *in vitro* and *ex vivo* studies

*Plasmodium falciparum* datasets are generated by synchronizing parasites chemically to specific stages on their developmental cycle within red blood cells. In the case of *P. vivax* datasets, blood from infected participants was added to culture. All these datasets were likely generated under constant environmental culture conditions, meaning that temperature, gas composition, and culture medium were held fixed throughout sampling, with no imposed light-dark, temperature, or feeding cycles, and no host-derived rhythmic signals (e.g., melatonin or body-temperature oscillations) supplied to the culture. These are the standard conditions for culturing *Plasmodium* parasites. Rhythms captured here therefore reflect endogenous parasite and host-cell oscillations driven by the IDC, rather than by external entrainment. This group includes multiple *P. falciparum* strains (3D7, FVO, SA250, D6, II3), *P. vivax ex vivo* IDC transcriptomes (9 datasets), and host-derived infection profiles (10 datasets).

### 3.5 Additional time-series studies or Repurposed non-circadian time-series

Although not explicitly designed for circadian studies, these datasets provide sufficient temporal sampling to infer secondary rhythms. Eight high-resolution *P. falciparum* transcriptomic datasets include IDC time courses (HB3, Dd2, 3D7), metabolic regulation experiments, and real-time transcription/mRNA decay measurements (3D7attB, 4-thiouracil labeling). We also identified 10 metabolomics datasets, profiling intra- and extracellular metabolite dynamics across synchronized IDC time courses in *P. falciparum* (3D7, NF54, and Dd2, including drug-perturbed conditions). These multi-omics datasets enable the detection of rhythms in transcription, protein abundance, and metabolism.

### 3.6 Distribution of omics data types

Transcriptomic datasets dominate the collection, comprising 82.08% (55 datasets), primarily using RNA-seq (40 datasets) and microarray (15 datasets) technologies (**Figure 3C**). Metabolomic datasets represent 14.92% (n = 10) and were obtained using mass spectrometry-based time-series profiling, while proteomic datasets comprise 3.0% (n = 2), including both gel-based and mass spectrometry-based proteome analyses.

### 3.7 Platform architecture, interface, and navigation

PlasmoRhythm is structured around a unified metadata framework that captures species, tissue, experimental conditions, sampling time points, and omics modality. Each transcript, protein, and metabolite is assigned a unique identifier and annotated using established community standards, including PlasmoDB gene names for parasites, DrugBank or STITCH identifiers for drug interactors, and Gene Ontology terms for virulence-associated genes. This harmonized structure supports reliable cross-study and cross-species comparisons.

All molecular entities link to interactive time-series visualizations that display raw and normalized expression profiles alongside fitted curves generated using four validated rhythmicity algorithms (JTK_CYCLE, ARSER, Lomb–Scargle, and meta2d). Each results page includes phase, period, amplitude, p-values, and q-values, all of which can be downloaded as a CSV file. Plots can be exported as JPG or PNG files. To maximize transparency and flexibility, PlasmoRhythm retains complete MetaCycle outputs for all analyzed features, allowing users to explore the data with different statistical cutoffs to evaluate the number of cycling entities in time-series datasets at varying levels of statistical stringency. Additional modules provide focused access to drug-interactor profiles and virulence-factor rhythmicity (**Figure 4).**

**Figure 4.**
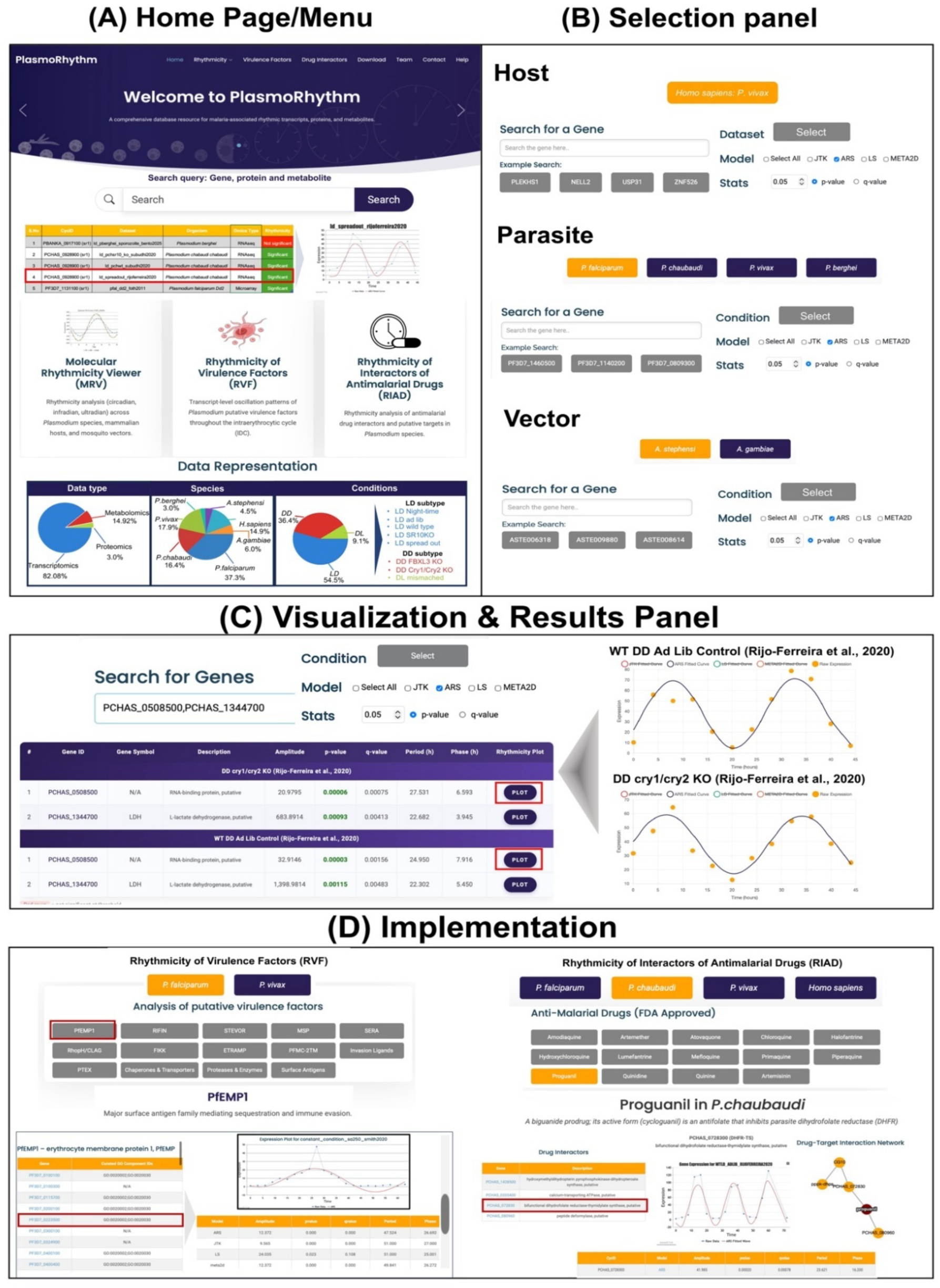
The web interface and key features of the PlasmoRhythm database. **(A)** Home Page/Menu. The landing page provides access to major modules, including molecular rhythmicity, rhythmic virulence factors, and rhythmic antimalarial drug interactors, along with interactive charts summarizing dataset distribution by species, omics type, and sampling strategy. **(B)** Selection Panel. Users can explore rhythmicity in Host, Parasite, or Vector modules, with advanced search and filter options by gene name, identifier, strain, condition, and rhythmicity detection model. **(C)** Visualization & Results Panel. Search results display oscillatory expression profiles for selected genes, proteins, or metabolites, with options to toggle between four rhythmicity evaluating algorithms (ARSER, JTK_CYCLE, Lomb-Scargle, meta2d). Plots show both model-fitted curves and raw data points, accompanied by downloadable rhythmic parameters. **(D)** Implementation. Systematically curated modules enable users to investigate the rhythmic expression of putative virulence factors (RVF module) and drug targets (RIAD module) associated with Plasmodium infections, providing interactive plots and downloadable data to support functional and translational research.

The PlasmoRhythm homepage provides a user-friendly entry point to all major database features (**Figure 4A**), offering interactive summaries of species, strains, conditions, and omics layers. A dedicated search panel enables queries by name, ID, or description, returning matched entries with organism details, functional annotation, omics type, and rhythmicity status (significant/nonsignificant). Selecting a result opens the corresponding detailed expression or abundance profile. This feature enhances transparency and usability, ensuring that users can easily navigate and understand the scope of PlasmoRhythm from the very first page.

A top navigation bar links to key modules: Home, Rhythmicity, Drug Interactors, Virulence Factors, Download, Team, Contact, and Help. The “Rhythmicity” module features dropdown filters to select a biological entity (Parasite, Host, or Vector) and an omics data type (Transcriptomics, Proteomics, or Metabolomics), enabling rapid exploration of time-series datasets across species and molecular layers. At the Download section of PlasmoRhythm, users can access three files per dataset: (i) meta2D output, (ii) the processed expression matrix generated after preprocessing, and (iii) the original data file. Dataset-specific preprocessing steps, filtering criteria, and MetaCycle settings are provided in Supplementary **Table S1**.

To enhance accessibility, the homepage also includes three shortcut panels that provide direct access to commonly used modules: Molecular Rhythmicity Viewer (MRV), Rhythmicity of Virulence Factors (RVF), and Rhythmicity of Interactors of Antimalarial Drugs (RIAD). These panels offer quick access to key features and facilitate navigation across the database. These shortcut panels complement the top navigation bar by providing direct access to the three principal analysis modules.

## 4. Modules and functions

### 4.1 Molecular Rhythmicity Viewer - MRV module

The MRV offers a direct entry point on the homepage/main menu to explore transcript-level time-dependent expression profiles across hosts, *Plasmodium* parasites, and mosquito vectors (**Figure 4A and 4B**). All panels share a uniform interface, ensuring a consistent user experience regardless of the selected organism. Users can search and filter genes by PlasmoDB ID, name, species, strain, or experimental condition, and compare results across multiple datasets. The search interface also enables users to query multiple candidates simultaneously across selected datasets and rhythmicity-detection algorithms. Rhythmicity parameters for all queried targets are presented in a unified table, with options to visualize individual expression profiles and download the results for further analysis. This feature enables users to compare the rhythmic patterns of multiple selected candidates simultaneously.

Results are presented as interactive plots that display raw expression data alongside rhythmic model fits **(Figure 4C**). Tables summarizing rhythm parameters, such as period, phase, amplitude, *p*-value, and *q*-value, are also provided. All outputs are downloadable for offline analysis or figure preparation. Users can also easily select different species, strains, models, or datasets based on their study goal, facilitated by an easily switchable top navigation bar **(Figure 4B)**.

Besides the MRV module, the Rhythmicity tab (in the top navigation bar) is the central interface for exploring temporal datasets across the malaria transmission system **(Figure 4B–D)**. Users can proceed with the analysis by selecting a biological entity (Parasite, Host, or Vector) and the desired omics layers (Transcriptomics, Proteomics, and Metabolomics). Each module is described in detail below.

### 4.2 Rhythmicity of Virulence Factors - RVF module

Host susceptibility to malaria has been shown to correlate with circadian immune function, metabolism, and hormone signaling (Rankawat *et al*. 2023, Hirako, Ramalho, and Gazzinelli 2025). The *Plasmodium* parasite is thought to exploit this rhythmic host environment by synchronizing its IDC with the host circadian clock (O’Donnell *et al*. 2011). This temporal alignment represents an adaptive strategy that maximizes parasite fitness, contributes to virulence, and facilitates recovery from temporal perturbations (Carvalho Cabral *et al*. 2024). Notably, the rhythmic expression of major virulence factors in malaria could also be a way for the parasite to exploit host rhythms to its advantage. For example, the presentation of *P. falciparum* Erythrocyte Membrane Protein 1 (PfEMP1), a key mediator of cytoadherence and immune evasion, is tightly constrained to the rhythmic IDC (Wahlgren, Goel, and Akhouri 2017, Rijo-Ferreira *et al*. 2020, Motta *et al*. 2023). Furthermore, while antigenic switching is an epigenetic process (Merrick *et al*. 2012). The IDC’s synchrony imposes a temporal gate on the expression of new gene variants, linking circadian timing to the parasite’s strategy for long-term persistence and severe disease (Recker *et al*. 2011) The RVF module of PlasmoRhythm provides access to curated rhythmic expression profiles of putative *Plasmodium* virulence genes, based on experimental literature and functional annotations. The gene families included are PfEMP1, RIFIN, STEVOR, MSP, SERA, CLAG, and FIKK kinases, all of which are involved in either immune evasion, erythrocyte invasion, or pathogenesis (Li Q *et al*. 2025).

To further characterize the functional landscape of these virulence factors, Gene Ontology (GO) enrichment analysis was performed on the gene sets from *P. falciparum and P. vivax*. Additionally, the corresponding GO terms and GO identifiers for each virulence gene are readily accessible in the database. The most enriched biological processes include immune evasion, host cell modification and adhesion, and include both shared and species-specific aspects of parasite virulence.

By implementing this RVF module, users can explore the temporal dynamics of the expression of these virulence genes and their families across multiple datasets. Rhythm parameters, including period, phase, and amplitude, are available for download. The interface supports comparisons across strains and experimental conditions, enabling users to investigate how gene expression timing varies in different contexts. This module supports investigations into the temporal regulation of virulence and studies on disease progression, host–parasite interactions, and chronotherapeutic strategies **(see Case Study 2, Figure 5D-E)** (Detailed case studies are discussed in the discussion section).

**Figure 5.**
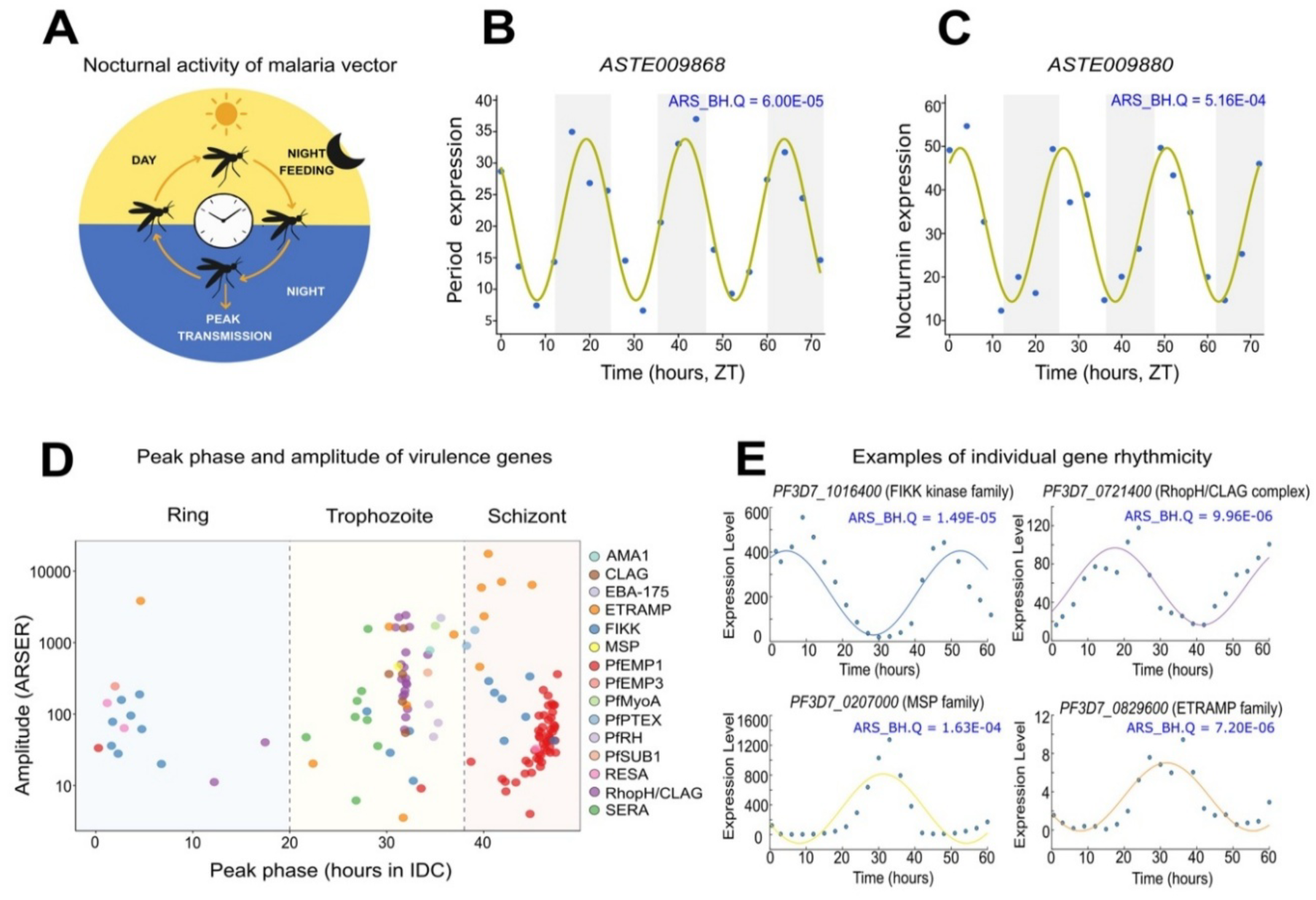
Examples of data access in PlasmoRhythm, demonstrating the analysis of representative rhythmic gene expression profiles in the malaria vector and parasite. **(A–C) Case study 1: PlasmoRhythm enables the visualization of the periodic expression of genes in the mosquito vector *A. stephensi*, facilitating an understanding of time-dependent behavioral modulation.** **(A)** Schematic illustrating the time-dependent activity pattern of the malaria vector, with peak feeding and transmission occurring at night. **(B, C)** Temporal expression profiles of two representative genes, *ASTE009868* (period expression, B) and *ASTE009880* (nocturnin expression, C), in *A. stephensi* under light-dark (LD) conditions. Both genes display robust 24-h rhythmicity, as determined by ARSER (yellow line: model fit; blue dots: raw data; ARS BH Q values indicated), with expression peaks aligned to nocturnal activity windows. **(D–E) Case study 2: Analysis of the rhythmic expression patterns of the major virulence factors across the intraerythrocytic developmental cycle (IDC) in *P. falciparum*. (D)** Phase-amplitude plot showing the distribution of peak phase (h in the IDC) and amplitude (ARSER) for prominent virulence gene families, including AMA1, CLAG, EBA-175, ETRAMP, FIKK, PfEMP1, MSP, RIFIN, and others. Genes cluster by IDC stage (ring, trophozoite, schizont), indicating coordinated temporal regulation. **(E)** Examples of individual rhythmic expression profiles for a few selected virulence genes: *PF3D7_1016400* (FIKK kinase family), *PF3D7_0721400* (RhopH/CLAG complex), *PF3D7_0207000* (MSP family), and *PF3D7_0826900* (ETRAMP family). Blue dots represent raw data; yellow lines show ARSER model fits.

**Figure 6.**
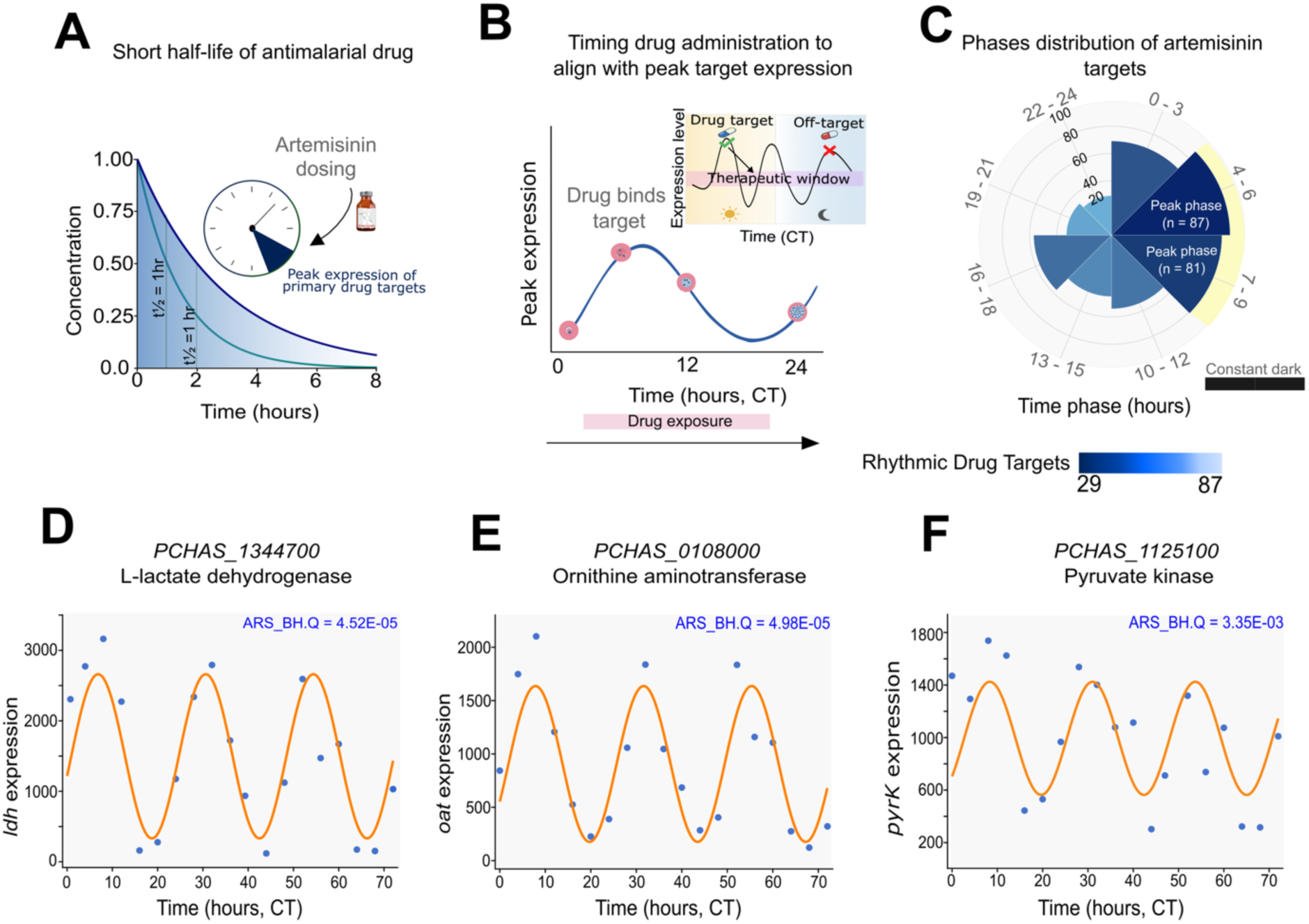
Evaluation of rhythmic expressions of antimalarial drug interactors in the PlasmoRhythm database (Case study 3). **(A)** Pharmacokinetic profile illustrating the short plasma half-life of the most effective antimalarial drugs, artemisinin and its derivatives, emphasizing the need for precise timing of administration for the possible improvement of their therapeutic efficacy. **(B)** Schematic depicting the concept of aligning drug dosing time with the peak expression of targets to maximize drug-target engagement and therapeutic efficacy. **(C)** Phase distribution of the rhythmic artemisinin targets in *P. chabaudi* under constant darkness, with the majority of targets peaking between circadian time (CT) 4–9 h. **(D–F)** Temporal expression profiles of three representative artemisinin targets, L-lactate dehydrogenase (*PCHAS_1344700*, D), ornithine aminotransferase (*PCHAS_0108000*, E), and pyruvate kinase (*PCHAS_1125100*, F), showing robust 24 h rhythmicity as determined by ARSER (orange line: model fit; blue dots: raw data; ARS_BH.Q values indicated). Together, these panels illustrate how PlasmoRhythm enables identification of optimal dosing windows based on the circadian regulation of antimalarial targets in the malaria parasites.

### 4.3 Rhythmicity of Interactors of Antimalarial Drugs - RIAD module

Canonical pharmacokinetic processes, including absorption, distribution, metabolism, and excretion (ADME), exhibit robust 24-h rhythms (Ruben *et al*. 2019, Dong *et al*. 2020). These rhythms are pronounced in the liver, where oscillating cytochrome P450 enzymes drive drug metabolism and where the malaria parasites first establish infection (Reuling *et al*. 2018, Chen *et al*. 2020). The goal of chronotherapy is to maximize on-target efficacy against the parasite while minimizing off-target toxicity in the host, thereby improving the therapeutic index (Dong *et al*. 2020, Lee, Field, and Sehgal 2021).

Evaluation of dosing-time dependency is especially critical for antimalarials with short half-lives, such as artemisinin derivatives (t₁/₂ = ∼1.5 h), which exhibit rapid fluctuations in plasma concentration (Siqueira-Neto *et al*. 2023). Thus, when exploring the optimal timing of treatment, it is important to consider rhythmic gene expression in both the parasite and the host (Cambie *et al*. 1991, Landau *et al*. 1992, Motta *et al*. 2023, Rankawat *et al*. 2023). In this context, optimizing treatment to a simple once-daily dosing schedule that aligns with host and parasite rhythms can enhance drug efficacy while minimizing the potential side effects and the financial and logistical burdens associated with multiple daily doses, thereby improving patient adherence to the therapy regimen and treatment outcomes.

The relevance of time-dependency on drug efficacy in antimalarial treatment has been proposed. For example, the evaluation of the dosing time dependency of artemisinin in the murine malaria model indicates that the drug is most effective if administered when the parasites are late rings or early trophozoites, as heme is required for drug activation (Owolabi, Reece, and Schneider 2021). Therefore, deploying effective chronotherapy requires a better understanding of when parasite drug targets are active.

To this end, PlasmoRhythm can support systematic analysis of the 24-h transcriptional rhythms of *Plasmodium* genes that encode known targets or interactors of antimalarial drugs. The RIAD module of PlasmoRhythm presents rhythmic expression profiles of *Plasmodium* genes predicted to interact with FDA-approved antimalarial drugs. Targets were curated from the STITCH database (Szklarczyk *et al*. 2016) and supplemented with manually annotated literature evidence. The RIAD interface displays the corresponding evidence source and confidence information for each drug–target interaction, enabling users to distinguish experimentally supported interactions from computationally inferred associations.

Users can explore how the expression of interactors of antimalarial drug dynamics varies across parasite species and IDC stages, aiding the identification of optimal drug administration windows. For each gene, rhythmic profiles are displayed as interactive time-series plots, accompanied by downloadable parameters (e.g., period, phase, amplitude, *p*-value, *q*-value). These features facilitate rhythm-based drug prioritization and the study of stage-specific drug sensitivity. By mapping the expression peaks of drug targets to specific IDC phases, the module supports research into therapeutic timing strategies and resistance mechanisms **(Case Study 3, Figure 6A– F)** (Detailed case studies are discussed in the discussion section).

## 5 DISCUSSIONS

Temporal regulation is crucial in malaria biology, particularly in coordinating the development of the parasite with that of the host and vector. In recent years, several in-depth, systems-level time-series studies have been reported on malaria parasites, hosts, and vectors (Rijo-Ferreira *et al*. 2020, Habtewold *et al*. 2022, Motta *et al*. 2023, Bento *et al*. 2025, O’Donnell, Prior, and Reece 2020). However, to date, there is no comprehensive, dedicated database available that supports the integration and analysis of these heterogeneous, large-scale time-series datasets. PlasmoRhythm addresses this gap by offering the first open-access, multi-omics web platform to document and evaluate the rhythmic expression of genes, proteins, and metabolites across the malaria transmission cycle.

Existing prominent database resources in circadian biology, such as CircaDB (Pizarro *et al*. 2013), CircadiOmics (Ceglia *et al*. 2018), CirGRDB (Li X *et al*. 2018), and CircaKB (Zhu *et al*. 2025), primarily focus on the circadian and diurnal landscape in mammals and other model organisms, but lack the rhythmicity datasets on malaria parasites. Although a few generic rhythmicity databases, such as RhythmicDB and CircadiOmics, include a limited number of datasets related to malaria, an integrated database focusing on the rhythms of multiple hosts, parasite species and vectors involved in this complex infection was missing.

In this regard, PlasmoRhythm supports interactive exploration of circadian (periodicity ∼24 h) and infradian (periodicity >24 h) rhythms in *P. falciparum, P. vivax, P. chabaudi, P. berghei*, and their mammalian and mosquito counterparts, under diverse experimental conditions including constant darkness, light/dark entrainment, and *in vitro* developmental cycles. It contains carefully curated datasets (many of which were published very recently) that are not available elsewhere. The comprehensive integration of these diverse, high-throughput omics-scale data sets makes PlasmoRhythm uniquely suited to fill a critical gap in malaria chronobiology, enabling new systems-level insights into host-parasite–vector temporal interactions.

PlasmoRhythm is not limited to evaluating molecular rhythmicity in malaria parasites, hosts, and vectors, but also includes two imperative modules (RVF and RIAD) to support the analysis of rhythmicity in malaria virulence factors and antimalarial drug interactors, thereby enhancing the translational importance of PlasmoRhythm.

The following representative case studies illustrate how PlasmoRhythm can be applied to translational and mechanistic analyses. An additional strength of PlasmoRhythm is its bulk-download facility, which provides access to original datasets, processed expression matrices, preprocessing and analysis metadata, and complete MetaCycle outputs. This feature promotes transparency and reproducibility while enabling independent reanalysis and integration with external computational workflows.

### 5.1 Analysis of time-dependent behavioral modulation in the *Anopheles* mosquito

PlasmoRhythm could be used to support vector biology research. We provide an example in Case Study 1 of analyzing transcriptional rhythms in *A. stephensi* under light-dark conditions, indicating diel oscillations in core circadian genes, such as period (ASTE009868; ARSER BH. Q = 6.00e-05, JTK BH. Q = 0.02, LS BH. Q = 0.9, meta2d_BH. Q = 7.6E-09) and nocturnin (ASTE009880; ARSER BH. Q = 5.16E-04, JTK BH. Q = 5.00e-02, LS BH. Q = 0.9, meta2d_BH. Q = 8.21e-07).

These gene expression patterns matched the mosquito’s nocturnal biting activity (Bento *et al*. 2025) **(Figure 5A–C)**. Understanding of 24-h molecular rhythms may help explain why mosquito transmission efficiency varies according to the time of day (Bento *et al*. 2025, Ochwedo *et al*. 2025, Stopard *et al*. 2025).

### 5.2 Rhythmicity in malaria virulence factors

PlasmoRhythm enables systematic analysis of the rhythmic expression of virulence factors in *P. falciparum*. In Case Study 2, we demonstrate how users can assess the temporal expression patterns of prominent virulence gene families, including PfEMP1, RIFIN, STEVOR, MSP, SERA, FIKK kinases, and ETRAMPs. These gene families exhibit clear oscillatory expression, with peaks during the late trophozoite and schizont stages, which are important for immune evasion and erythrocyte invasion (Smith *et al*. 2020) **(Figure 5D–E)**. Representative rhythmic genes included *PF3D7_1016400* (FIKK kinase family; ARSER BH.Q = 1.49e-05, JTK BH.Q = 7.17e-04, LS BH.Q = 1.05e-01, meta2d BH.Q = 3.20e-08), *PF3D7_0721400* (RhopH/CLAG complex; ARSER BH.Q = 9.96e-06, JTK BH.Q = 1.24E-04, LS BH.Q = 1.03e-01, meta2d BH.Q = 4.92E-11), *PF3D7_0207000* (MSP family; ARSER BH.Q = 1.63e-04, JTK BH.Q = 5.53e-04, LS BH.Q = 1.16e-01, meta2d BH.Q = 3.79e-07), and *PF3D7_0826900* (ETRAMP family; ARSER BH.Q = 9.07e-05, JTK BH.Q = 9.41e-04, LS BH.Q = 1.03e-01, meta2d BH.Q = 2.08e-07. The rhythmic expression of malaria virulence factors underscores the need to account for these regulatory dynamics when designing vaccines and interventions targeting antigens expressed during these key IDC phases.

### 5.3 Evaluation of the dosing-time dependency of antimalarial drugs

PlasmoRhythm extends beyond exploratory chronobiology to provide practical tools for translational research, including optimizing antimalarial drug efficacy through chronotherapy. In Case Study 3, we demonstrated how users can analyze the rhythmic expression of artemisinin targets in *P. chabaudi* in constant dark conditions (Rijo-Ferreira *et al*. 2020). We selected artemisinin and its derivatives for this analysis because they have a short half-life in human plasma, typically ranging from 1.5-5 h (Tilley *et al*. 2016). Many targets of artemisinin and its derivatives, such as lactate dehydrogenase (*PCHAS_1344700;* ARSER BH. Q = 4.52e-05, JTK BH.Q = 5.44e-04, LS BH.Q = 0.13, meta2d_BH.Q = 1.04e-08), ornithine aminotransferase (*PCHAS_0108000;* ARSER BH. Q = 4.98e-05, JTK BH.Q = 4.64e-04, LS BH. Q = 0.13, meta2d_BH.Q = 8.28e-09) and pyruvate kinase (*PCHAS_1125100;* ARSER BH. Q = 3.35E-03, JTK BH.Q = 8.56E-03, LS BH.Q = 2.25e-01, meta2d_BH.Q = 1.63e-04), exhibited peaks between circadian time (CT) 4 and 9 h **(Figure 6A–F)** and exhibited overall statistically significant rhythms, as determined by ARSER with Benjamini-Hochberg correction (*q* < 0.05) (Rijo-Ferreira *et al*. 2020). These findings highlight the potential for chronotherapy, whereby short-half-life antimalarials, such as artemisinin, may be timed to coincide with peak target abundance to improve efficacy (Cambie *et al*. 1991; Owolabi, Reece, and Schneider 2021).

## 6 Conclusion and future directions

As malaria research evolves toward more integrated approaches, an easy-to-use and comprehensive web platform like PlasmoRhythm could play a crucial role in this evolution. The platform helps identify the optimal timing of treatments, determine when parasites are most susceptible to various drugs, and explore how biological rhythms influence malaria transmission and immune responses, thereby informing the development of new strategies for more effective malaria control.

Although several transcriptomic time-series datasets have emerged in recent years in malaria research, encompassing parasites, hosts, and vectors, very few temporally resolved, high-resolution proteomic or metabolomic profiles are available. Notably, there is a dearth of temporally resolved midgut omics datasets from mosquito vectors, despite the midgut being a key site of parasite development and transmission. Also, there is a paucity of omics-level time-series datasets from malaria parasites derived from clinical or field isolates. We are optimistic that our comprehensive repository, PlasmoRhythm, will be further enhanced with the arrival of additional omics-level high-resolution datasets, particularly at the post-transcriptional and post-translational levels.

The current rhythmicity-detecting algorithms used in PlasmoRhythm (JTK_CYCLE, ARSER, Lomb–Scargle, MetaCycle/meta2d) assume stationarity, including stable period, phase, and amplitude, which may not capture dynamic changes in host–parasite interactions, where circadian cues, feeding schedules, environmental conditions, or pharmacological interventions can influence developmental timing. In this regard, Continuous Wavelet Transform (CWT)-based approaches, such as pyBOAT (Schmal, Mönke, and Granada 2022), may provide complementary time-frequency analyses that estimate instantaneous changes in rhythmic parameters without requiring stationarity assumptions, which could be a potential future inclusion for our database. However, robust wavelet analysis requires densely sampled, regularly spaced data, which many existing time-series datasets in malaria biology lack (particularly at the proteome and metabolome levels). As higher-resolution datasets become increasingly available, incorporating wavelet-based frameworks will enhance PlasmoRhythm’s ability to characterize dynamic rhythmic processes across host, parasite, and vector systems. Community submission tools and interoperability between existing malaria databases will be introduced to support sustainability, enabling PlasmoRhythm to function as a continuously updated, community-driven resource. All in all, as interest grows in using biological timing to improve health outcomes, PlasmoRhythm provides an essential resource for diverse research communities investigating malaria biology, chronobiology, drug development, and disease prevention.

## ACKNOWLEDGMENTS

The authors thank all researchers who generated and publicly released the data included in our PlasmoRhythm database. The authors thank all the members of the labs for discussions on key datasets and how best to present the data, and acknowledge computational infrastructure support from the Indian Institute of Technology Hyderabad for hosting the database.

## AUTHORS’ CONTRIBUTIONS

S. Ray, R.K., and S. Rankawat conceptualized the database. S. Rankawat, K.K., S.S., S.D., B.P., B.T.K., F.R.F., R.K., and S. Ray performed the data analyses and result presentation. S. Ray, R.K., and F.R.F. supervised dataset curation. K.K., S. Rankawat, S.S., B.T.K., and R.K. constructed the database. S. Rankawat and S. Ray wrote the manuscript and prepared the figures. All authors contributed to editing and reviewing the final manuscript.

## DATA AVAILABILITY

All datasets used in this study are publicly available and detailed in the Supplementary Material. These datasets can also be downloaded from the PlasmoRhythm Download section. PlasmoRhythm version 1.0, released in February 2026, is freely accessible without registration at https://project.iith.ac.in/cgntlab/PlasmoRhythm/.

## SUPPLEMENTARY DATA FILES

**Table S1. Summary of omics datasets included in PlasmoRhythm.** Comprehensive listing of all time-series omics datasets integrated into PlasmoRhythm, including organism, platform, tissue, experimental conditions, number of time points, sample size, temporal resolution, rhythmicity analysis algorithms used, data type, availability, and references for each dataset.

